# Genomic Islands in *Wolbachia* Prophages Drive Amplification and Diversification of Cytoplasmic Incompatibility Genes in *Culex pipiens*

**DOI:** 10.1101/2025.10.03.680192

**Authors:** Julien Amoros, Alice Namias, Théo Durand, Haoues Alout, Julien Martinez, Mylène Weill, Mathieu Sicard

**Affiliations:** ISEM, Université de Montpellier, CNRS, IRD, EPHE, Montpellier, France; MIVEGEC, Université de Montpellier, CNRS, IRD, Montpellier, France; Université Paris-Saclay, CNRS, AgroParisTech, Ecologie Société Evolution, 91190, Gif-sur-Yvette, France; ASTRE, Université de Montpellier, INRAe, CIRAD, Montpellier, France; Tree of Life, Wellcome Sanger Institute, Hinxton, United Kingdom

## Abstract

*Wolbachia* are maternally inherited endosymbiotic bacteria widespread among arthropods. They manipulate their host reproduction to enhance their prevalence in host populations. The most common manipulation is cytoplasmic incompatibility (CI), causing embryonic death in crosses between infected males and uninfected females, or between individuals carrying distinct incompatible *Wolbachia*. CI patterns are highly complex in the mosquito *Culex pipiens*, where the causal genes *cidA* and *cidB* are amplified and diversified, forming a “*cid* repertoire” within each *Wolbachia w*Pip genome. Despite their central role in CI, the genomic mechanisms underlying such *cid* amplification and diversification remained poorly understood. This knowledge gap is largely due to the difficulty of assembling *w*Pip’s genomes due to highly repeated genes and mobile elements, especially in WO prophages. Here, we directly annotated Illumina polished Nanopore-sequences to investigate the genomic flanking context of *cid* genes in three distinct *w*Pip lineages. We assembled WO prophage regions of substantial length containing the entire *cid* repertoire previously described in these bacterial lineages. Within these WO regions, *cid* genes are consistently embedded in modular and rearrangeable islands composed of *MutL, rnhA*, and small mobile elements, all displaying hyperconserved nucleotide identity across islands. These genomic islands are probably drivers for major rearrangement and recombination events responsible for the amplification and diversification of *cid^w^*^Pip^ genes within and between the *w*Pip genomes leading to CI complexity in *C. pipiens*.

## Introduction

*Wolbachia* are highly prevalent endosymbiotic bacteria infecting arthropods and nematodes, responsible for the largest known pandemic among metazoans (Zug and Hammerstein 2012; Weinert et al. 2015). The best-known *Wolbachia*-phenotype is Cytoplasmic Incompatibility (CI), a form of reproductive parasitism that reduces egg hatching rate when *Wolbachia* infected males mate with uninfected females, or between individuals carrying incompatible *Wolbachia* strains (Yen and Barr 1973; Breeuwer and Werren 1990; O’Neill and Karr 1990; Bordenstein et al. 2001; Duron et al. 2006; Atyame et al. 2014). CI is primarily induced by CI Inducing Factor (Cif) proteins encoded by *cif*A-*cif*B genes arranged in tandems inside *Wolbachia* genomes (LePage et al. 2017; Beckmann et al. 2017; Lindsey et al. 2018). To date, ten phylogenetic groups of *cif* genes have been identified, named *cif* types I - X (Lindsey et al. 2018; Martinez et al. 2021; Tan et al. 2024; Amoros et al. 2025). Among them, type I (from *w*Mel and *w*Pip), type III (from *w*No) and type IV (from *w*Pip) were functionally proven to induce and rescue CI through transgenic expression in *Drosophila* (LePage et al. 2017; Beckmann et al. 2017; Chen et al. 2019; Sun et al. 2022). These genes can also be classified according to their functional domains: *cif* type I genes containing an active deubiquitinase (DUB) domain are functionally classified as *cid* genes (for CI-inducing DUB) while *cif* types II, III and IV genes, containing active nuclease domains (Martinez et al. 2021), are functionally classified as *cin* (CI-inducing nuclease). Transgenesis experiments in *Drosophila* and yeasts cells demonstrated that the B genes encode toxins preventing the normal cell culture growth, while the A genes encode antidotes which restore normal growth (Beckmann et al. 2017; Chen et al. 2019; Sun et al. 2022; Horard et al. 2022; Terretaz et al. 2023; Namias et al. 2025).

In the mosquitoes of the *Culex pipiens* complex, in which >99% individuals are infected with *w*Pip *Wolbachia* (Klasson et al. 2008; Dumas et al. 2013), crossing between dozens of lineages revealed highly complex CI patterns, including both unidirectional and bidirectional incompatibilities (Laven 1967; Duron et al. 2006; Atyame et al. 2014) (Supplementary Table 1). A Multiple Loci Sequencing Typing (MLST) approach, specific to *w*Pip, identified five distinct clades named group *w*Pip-I to *w*Pip-V (Atyame et al. 2011; Dumas et al. 2013) (fig 1A), which strongly correlate with CI patterns. Indeed, mosquitoes infected with *w*Pip from the same group are generally compatible, while compatibility in crosses between individuals infected with distinct groups is less predictable (Atyame et al. 2014). These results, along with backcross experiments demonstrated that genetic variations in *w*Pip, but not in the host genetic background, are responsible for CI patterns complexity in *C. pipiens* (Duron et al. 2005; Atyame et al. 2011; Atyame et al. 2014; Sicard et al. 2021). Before the discovery of *cif* genes, both theoretical and experimental studies had concluded that each *w*Pip genome must carry multiple and polymorphic genes responsible for CI to explain complex CI patterns in *C. pipiens* (Nor et al. 2013; Atyame et al. 2014). Since then, *cid^w^*^Pip^ (*i.e*. type I *cif*) genes were indeed found to be amplified and diversified in all *w*Pip genomes studied and are now recognized as the main drivers of the *Culex*-CI complexity (Bonneau et al. 2018a; Bonneau et al. 2019; Sicard et al. 2021; Namias et al. 2024a; Namias et al. 2025) (Table 1). Each *w*Pip genome carries up to six distinct *cidA-B* pairs, with two forms of *cidB^w^*^Pip^ co-occurring: (i) a full length *cidB^w^*^Pip^ variant of around 3.5kb, bearing a DUB domain and demonstrated as key for CI phenotypes (Beckmann et al. 2017; Beckmann et al. 2019a; Horard et al. 2022; Namias et al. 2025), and (ii) several truncated *cidB^w^*^Pip^ variants, lacking part or all of the DUB domain that are less stable (Namias et al. 2025). This arsenal of *cid^w^*^Pip^ genes, composed of distinct *cidA^w^*^Pip^ and *cidB^w^*^Pip^ variants in one genome, was called a “*cid* repertoire” (Bonneau et al. 2018a; Namias et al. 2024a; Namias et al. 2025). In addition to these *cidA-B^w^*^Pip^ pairs, all the *w*Pip genomes studied so far present *cif* genes from the type IV, called *cinA-B^w^*^Pip^. As these *cin* genes are present in a single copy and exhibit no polymorphism between *Culex* lines, their implication in CI complexity in *Culex* was ruled out (Bonneau, Atyame, et al. 2018; Martinez et al. 2021) (Table 1).

**Table 1.** Overview of previous data obtained on *cif* genes in the *w*Pip genomes analyzed in this study. ^A^ assembled draft genomes (Klasson et al. 2008; Salzberg et al. 2009; Pinto et al. 2013); ^B^ number quantified through qPCR (taking *wsp* monocopy gene as reference; (Bonneau et al. 2018b); ^C^ diversity assessed using Nanopore sequencing of *cidA* and *cidB* PCR amplicons on single individual (Namias et al. 2025); ^D^ Analysis of depth of coverage and polymorphism by mapping *cinA/cinB* Illumina reads on *w*Pip-Pel genome (Bonneau et al. 2018a); - missing information, ∼ partial information; *number depending on the individual analysed

| **wPip name**<br>Accession number | **wPip group** | **Mosquito host species** | **Geographical**<br>**origin of host**<br>***Culex***<br>**mosquito** | **Mosquito**<br>**lineage type** | **Estimated**<br>**number of**<br>***cidA/cidB***<br>**operon per**<br>**genome** | **Estimated**<br>**number of**<br>**different**<br>***cidA/cidB***<br>**variants per**<br>**genome** | **Estimated**<br>**number of**<br>***cinA/cinB***<br>**operon per**<br>**genome** | **Estimated**<br>**number of**<br>**different**<br>***cinA/cinB* variants**<br>**per genome** |
| --- | --- | --- | --- | --- | --- | --- | --- | --- |
| Pel<br>AM999887.1 | wPip-I | *Culex pipiens*<br>*quinquefasciatus* | Sri Lanka | Colony | 1/ 1<sup>A</sup> | 1/ 1<sup>A</sup> | 1/ 1<sup>A</sup> | 1/ 1<sup>A</sup> |
| JHB<br>NZ_ABZA00000000.1 | wPip-I | *Culex pipiens*<br>*quinquefasciatus* | South Africa | Unknown | 2/ ~3<sup>A</sup> | 2/~3<sup>A</sup> | 1/ 1<sup>A</sup> | 1/ 1<sup>A</sup> |
| Tunis | wPip-I | *Culex pipiens pipiens* | Tunisia | Isofemale<br>line | 6/ 6<sup>B</sup> | 3-4/ 3-5<sup>C*</sup> | 1/1<sup>D</sup> | 1/ 1<sup>D</sup> |
| Slab | wPip-III | *Culex pipiens*<br>*quinquefasciatus* | United States | Isofemale<br>line | 5/ 5<sup>B</sup> | 3-4/ 2<sup>C*</sup> | - | - |
| Mol<br>NZ_CTEH00000000.1 | wPip-IV | *Culex pipiens molestus* | China | Unknown | 0/ 0<sup>A</sup> | 0/ 0<sup>A</sup> | 1/ 1<sup>A</sup> | 1/ 1<sup>A</sup> |
| Harash | wPip-IV | *Culex pipiens pipiens* | Algeria | Isofemale<br>line | - | 4/ 3<sup>C</sup> | - | 1/ 1<sup>D</sup> |

Altogether, these findings significantly advanced our understanding on genetic determinism of unprecedented CI complexity induced by *w*Pip in *Culex*. However, the diversity of available *w*Pip genomes is currently too scarce to allow the study of genomic drivers that could explain such unprecedented diversification of *cid* genes between and within *w*Pip groups. To date, three *w*Pip draft genomes are currently available on public databases (*w*Pip-Pel, JHB and Mol) (Klasson et al. 2008; Salzberg et al. 2009; Pinto et al. 2013), Pel being the most extensively studied and the currently accepted reference genome, since it is made of a single non-circularized contig, and Mol the most incomplete and fragmented one (fig 1A). Since both *w*Pip-Pel and JHB belong to *w*Pip-I group (fig 1A), these genomes only capture a limited part of *w*Pip group diversity. Moreover, *cid^w^*^Pip^ gene copy numbers and diversity reported in these genomes do not match recent genetic datasets acquired on a broad range of *w*Pip (Bonneau et al. 2018a; Namias et al. 2023; Namias et al. 2024a; Namias et al. 2025) (Table 1). Indeed, the *w*Pip-Pel genome only contains a full length *cidB^w^*^Pip^ copy and no truncated ones while *w*Pip JHB has only two truncated *cidB^w^*^Pip^ copies (fig 1B). Strikingly, *cid^w^*^Pip^ genes are entirely absent from *w*Pip Mol contigs (fig 1B). This apparent discrepancy between recent *cif^w^*^Pip^ genetic data (Bonneau et al. 2018a; Namias et al. 2023; Namias et al. 2024a; Namias et al. 2025) and the three *w*Pip draft genomes (Klasson et al. 2008; Salzberg et al. 2009; Pinto et al. 2013) can be explained by the fact that none of these three genomes are fully circularized. This difficulty to robustly assemble *w*Pip genomes results from the highly repetitive nature of WO prophage genes (Bordenstein and Wernegreen 2004; Bordenstein and Bordenstein 2022; Vancaester and Blaxter 2023), which represent 20% of *w*PipPel and *w*PipJHB genomes (Klasson et al. 2008; Salzberg et al. 2009) (fig 1 B, C).

**Figure 1.**
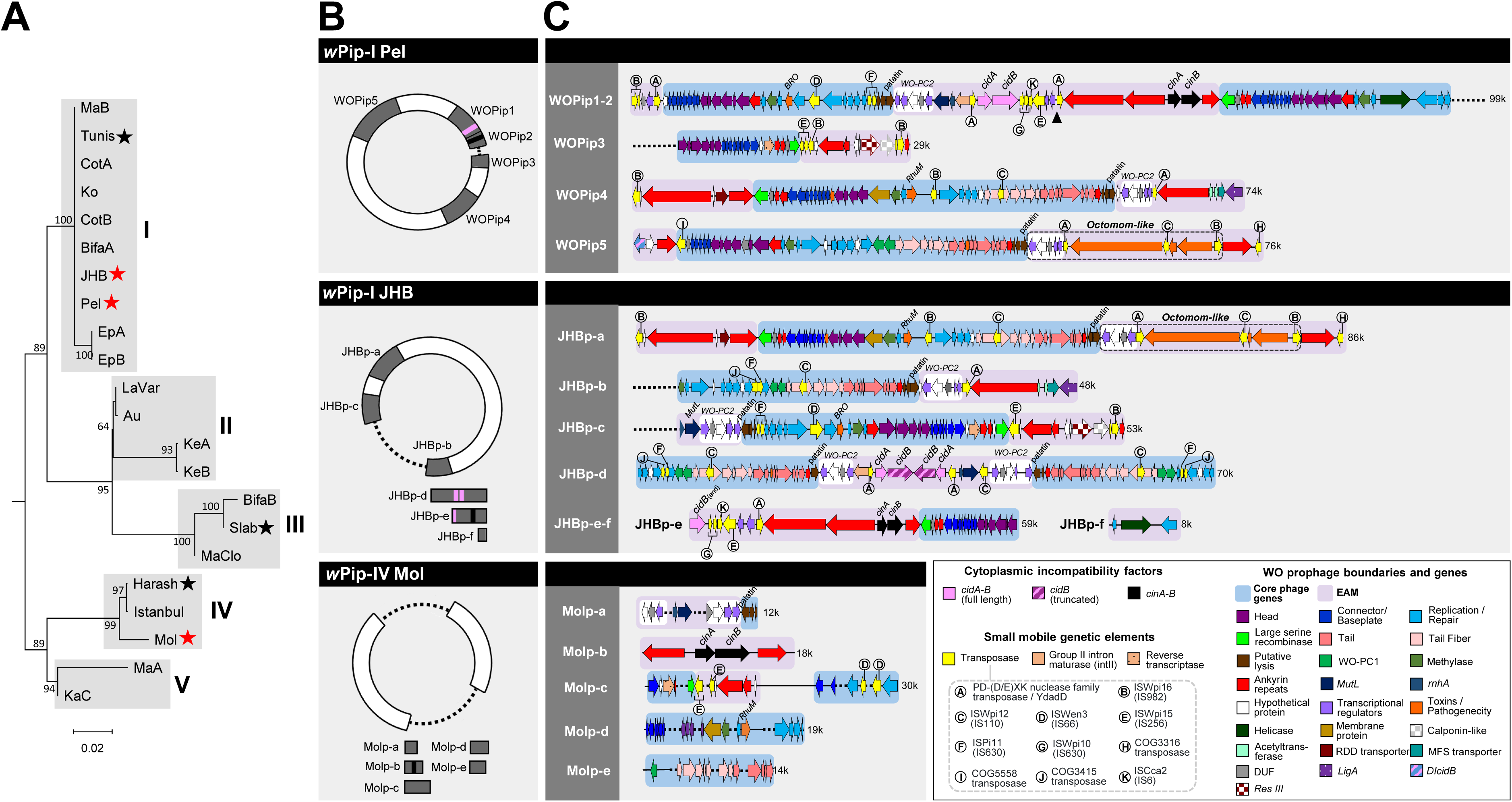
WO prophages from publicly available *w*Pip genomes (*w*Pip-Pel, JHB and *w*Pip Mol) (A) Maximum likelihood phylogeny of the five *w*Pip groups (I to V) based on five genes (*pk1, pk2, MutL, GP12*, and *GP15*) using 1,000 replicates (HKY+I+G4 substitution model, 4,012 nucleotides). Only bootstrap values greater than 60 are shown. Red stars indicate the three available public *w*Pip draft genomes, while black stars represent the *w*Pip genomic data analyzed in this study. (B) Genomic location of WO prophages and *cif* pairs (*cid* and *cin*) on the *w*Pip-Pel, JHB and *w*Pip-Mol chromosomes. The dark grey areas correspond to WO prophages, with the *cid* pairs highlighted in pink, and the *cin* pairs in black. The dotted lines indicate assembly gaps. For *w*Pip-JHB and *w*Pip-Mol, WO prophages are located in unassembled contigs. (C) Detailed (re)annotations of the WO prophages in the three public *w*Pip genomes. The conventional separation between WOPip1 and WOPip2 is marked by a black arrowhead as these two prophages occur sequentially in the chromosome. *DIcidB* refers to the isolated *cidB* fragments containing a deubiquitinase domain in *w*Pip-Pel (*WP1290–1291).* The length of each sequence in kb is indicated at the end of each prophage region.

WO prophages are basically composed of both core phage genes and Eukaryotic Association Module (EAM) genes (Bordenstein and Bordenstein 2016; Bordenstein and Bordenstein 2022) (fig 1 B, C). They are proposed to act as genomic hotspots that facilitate horizontal gene transfers between *Wolbachia* (Bordenstein and Wernegreen 2004; Kent et al. 2011; Bordenstein and Bordenstein 2016; Bordenstein and Bordenstein 2022). In the three available *w*Pip draft genomes, both *cid^w^*^Pip^ and *cin^w^*^Pip^ genes are located within WO prophage regions (Bordenstein and Bordenstein 2022) (fig 1C). If such locations suggest that WO prophages might be involved in *cid^w^*^Pip^ gene extreme amplification and diversification within and between *w*Pip genomes, the genomic mechanism(s) at play is not yet understood. A first putative mechanism is phage transduction, often considered the most straightforward explanation of horizontal transfer between *Wolbachia* genomes (Bordenstein and Reznikoff 2005; Kent et al. 2011; Bordenstein and Bordenstein 2016). While phage particles could theoretically promote *cid^w^*^Pip^ transfer between *w*Pip genomes, to date, the genome in virus particles attributed to WO phages have only been sequenced in *w*VitA-infected *Nasonia giraulti* and *w*Soc-infected *Allonemobius socius* and none of their genomes contained the *cif* genes (Bordenstein and Bordenstein 2016; Kupritz et al. 2021). An alternative mechanism to transduction is that the many gene repeats in WO prophages affect *cid* gene copy number and diversity by recombination and rearrangement during DNA replication (Slack et al. 2006; Lovett 2017). In addition, small mobile genetic elements, such as transposons and retrotransposons, within WO prophages have also been pointed out as major sources of *Wolbachia* genomic diversification (Cerveau et al. 2011; Leclercq et al. 2011; Cooper et al. 2019; Tan et al. 2024).

Here, by directly annotating Illumina polished Nanopore-sequences, we investigated the genomic environments of *cif^w^*^Pip^ genes (both *cid^w^*^Pip^ and *cin^w^*^Pip^) and the diversity of these environments among three different *w*Pip from three distinct groups, known to induce distinct CI patterns in their mosquito hosts: *w*Pip-I Tunis, *w*Pip-III Slab, and *w*Pip-IV Harash (Atyame et al. 2014; Sicard et al. 2021) (Supplementary Table 1). We found that *cid^w^*^Pip^ genes reside in modular and rearrangeable genomic islands within highly variable WO prophages, while the *cin^w^*^Pip^ genes are found in WO syntenic regions among the different *w*Pip. The *cid^w^*^Pip^ genomic islands harbor multicopy genes involved in horizontal transfer, recombination, and DNA repair which display hyperconserved nucleotide identity. These genomic islands are probably drivers for major rearrangement and recombination events responsible for the amplification and diversification of *cid^w^*^Pip^ genes within and between the *w*Pip genomes leading to CI complexity in *C. pipiens*.

## Materials and Methods

### Mosquito lines for Nanopore long read sequencing

Three mosquito isofemale lines, Tunis, Slab and Harash were used in this study (Table 1). Each isofemale line was founded with the progeny of a single egg raft from a single female. Mosquitoes were reared in 65 dm³ screened cages housed in a single room maintained at 25 °C with 70% relative humidity and a 12:12 light-dark cycle. Larvae were fed with a mixture of shrimp powder and rabbit pellets, while adults were provided with a honey solution. Females were blood-fed on turkey blood using a Hemotek membrane feeding system (Discovery Workshops, UK) to allow egg production. Each mosquito line is infected with a single *w*Pip genotype, previously characterized by specific MLST (Atyame et al. 2011; Dumas et al. 2013; Atyame et al. 2014): Tunis (*w*Pip-I), Slab (*w*Pip-III), and Harash (*w*Pip-IV) (fig 1A). Every three months, to check for contamination, a pool of a hundred larvae per line was collected, the DNA extracted, and the *w*Pip group determined using PK1 marker, as described in (Namias et al. 2024b).

High molecular weight genomic DNA was extracted from 10 to 20 females for each isofemale line. For Harash and Tunis lines, DNA was previously extracted using the Qiagen Genomic-tip 20G kit following the manufacturer’s protocol for insects (Namias et al. 2025) while for the Slab isofemale line, DNA was extracted using the Phenol-Chloroform extraction protocol with no vortexing to avoid DNA shearing. All DNA libraries were prepared using the Ligation Sequencing Kit (SQK-LSK109) and Native Barcoding Expansion Kit (EXP-NBD104) and then sequenced on a Minion Mk1B using a FLO-MIN106D flow cell with R9.4.1 chemistry. Short read DNA libraries were prepared in accordance with the Illumina DNA Prep Library guide and sequenced in NextSeq 550 Flowcell using the Illumina chemistry V2, 2×150bp paired end run. Trimmed long reads were then mapped to the *w*Pip-Pel reference genome to retrieve *Wolbachia* sequences with Minimap2 v2.29 (Li 2018). The long read sequences were polished with Illumina reads (2 × 150 bp) by using Pilon v1.24 (Walker et al. 2014) or Racon v1.4.20 and Ratatosk v0.9.0 (Vaser et al. 2017; Holley et al. 2021).

### *Wolbachia* sequences with *cif* genes and identification of *cidA-cidB* variants

The *Wolbachia* sequences (Nanopore corrected with Illumina) containing *cif^w^*^Pip^ genes (*cid* or *cin*) were sorted out by mapping corrected reads respectively on *cid* and *cin* sequences from *w*Pip-Pel reference genomes using Minimap2 v2.29 (Li 2018). Then, only sequences ≥10 kb containing *cif* genes were further analysed. *Cif* gene types were determined using the methodology described in Amoros et al. 2025. In sequences above 10 kb containing the *cid* genes, the *cidA* and *cidB* variants were characterized based on the *cid* references and nomenclature established by Namias et al. 2023; Namias et al. 2025. To achieve this, sequences ≥10 kb were independently aligned to *cidA* and *cidB* sequences from Namias et al. 2025 using MAFFT v7.505 (Katoh and Standley 2013).

### Annotations and assembly of *cif* gene environments in *w*Pip

Annotations were performed using BLASTn (megablast program) against the *w*Pip-Pel draft genome (accession number: AM999887.1) using the current version of the NCBI Prokaryotic Genome Annotation Pipeline (PGAP). Only nucleotide identity above 75% with *w*Pip-Pel was selected. Gene functions obtained from *w*Pip-Pel genome were then updated according to the new WO prophage gene annotations proposed by Bordenstein and Bordenstein 2022. To classify transposases, we conducted BLAST searches against ISfinder (https://isfinder.biotoul.fr/blast.php; Siguier et al. 2006) and ran HHpred (Zimmermann et al. 2018) with the SCOPe70 v.2.08, Pfam-A v37.0, SMART v6.0, and COG/KOG v1.0 databases. Group II intron retrotransposons (maturases) were identified using Zbase (http://webapps2.ucalgary.ca/~groupii/cgi-bin/main/blastusr.php, Candales et al. 2012). Then, the sequences sharing identical *cidA-cid* variants and exhibiting same annotated gene composition (with ≥99% nucleotide sequence identity), were aligned using MAFFT and manually assembled to form contigs (Supplementary Figure 1).

### Comparison of new WO prophage regions with reference ones

The gene compositions of newly identified WO prophages in draft genomes JHB and Mol, as well as those carrying *cif* genes in Tunis, Slab, and Harash, were compared to the gene compositions of the five reference prophages of *w*Pip-Pel (WOPip1 to WOPip5; fig. 1C). Each reference prophage displays distinctive specific genes that enable their identification: (i) WOPip1 is characterized by the BRO toxin (annotated as *WP0260*), replication/repair genes located near an IS630 and patatin-like putative lysis genes (fig 1C), (ii) WOPip2 by long genes containing ankyrin repeats and a helicase gene (*WP0319*), and (iii) WOPip3 is the only one containing both *ResIII* and calponin-like genes (*WP0350* and *WP0351*). Both WOPip4 and WOPip5 share structural modules such as tail and tail fiber genes, but differ in accessory content: WOPip4 presents an RDD transporter (*WP0409*), a membrane protein (*WP0428*), a RhuM toxin (*WP0431*), an acetyltransferase (*WP0463-0464*), and a *LigA* gene (*WP0465*), whereas WOPip5 harbors a COG5558 transposase (*WP1294*), WO-PC1 genes (*WP1319-1320*), and the octomom-like module (*WP1341-1350*). The presence/absence of these specific genes, as well as full gene composition of the reference prophages, were used to characterize the newly obtained WO prophage regions.

### Phylogenetic analyses

All phylogenetic analyses in this study were conducted using the maximum likelihood (ML) method with 1,000 bootstrap replicates using raxml-ng v1.1.0 (Stamatakis 2014). The best-fit evolutionary model was selected using modeltest-ng v0.1.7 (Darriba et al. 2020) based on the corrected Akaike Information Criterion (AICc). All network analyses were generated using the Neighbour-net method implemented in SplitsTree v4.19.2 (Huson and Bryant 2024). Sequence alignments were carried out using MAFFT v7.505 (Katoh and Standley 2013), implemented in Unipro UGENE v52.0 (Okonechnikov et al. 2012). The *w*Pip phylogeny was inferred from five multilocus genes: *pk1*, *pk2*, *MutL*, *GP12*, and *GP15* (fig1A) (Atyame et al. 2011). To identify the srWO groups of *w*Pip prophages, phylogenetic and networking analyses based on the amino acid sequences of the large serine recombinase were performed as defined by (Bordenstein and Bordenstein 2022). Additionally, seven widely distributed core WO prophage proteins, including structural, replication, and modification functions, were analysed by phylogenetic and network analyses using the same methods as described above. These included the Minor capsid protein (*WP0252*), Terminase large subunit GpA (*WP0255*), Tail tape measure protein (*WP0448*), Tail fiber protein (*WP1325*), Baseplate protein GpV (*WP0246*), RepA (*WP0433*), and Methylase (*WP0258*).

## Results

### The contigs with *cif* genes represent up to 20% of full *w*Pip genome

As previously reported in Namias et al. 2025, the Nanopore long reads polished with Illumina short reads obtained from both *w*Pip-I Tunis and *w*Pip-IV Harash lines, did not allow the *de novo* assembly of significant parts of respective *Wolbachia* genomes. For *w*Pip-III Slab, newly analysed here, we obtained a total number of 4,886 long reads (with N50=1.3kb). The mapping to the reference *w*Pip-Pel genome showed a coverage of 98.8% and a depth of coverage of only 8X. This poor depth of coverage for *w*Pip-III Slab also prevented us from obtaining a full genome *de novo* assembly. However, sequences ≥10 kb harbouring *cif* genes were recovered from the whole-genome datasets (*ie*. long reads corrected with illumina) of *w*Pip-I Tunis, *w*Pip-III Slab and *w*Pip-IV Harash (Supplementary Table 2). Individual annotations followed by alignment of similar annotated sequences allowed us to manually assemble contigs harbouring *cif* genes for each *w*Pip line. Among them, we reconstructed four different contigs for *w*Pip-I Tunis (named Tunisp-a to d, p for prophage), three different contigs for *w*Pip-III Slab (named Slabp-a to c), and four different contigs for *w*Pip-IV Harash (named Harashp-a to d) (Supplementary Figure 1; Supplementary Table 2). Altogether, the contigs represent approximately 200 kb for Tunis, 154 kb for Slab, and 300 kb for Harash, corresponding to 13-20% of *w*Pip genome, which is estimated to be ∼1.5 Mbp (fig 2).

**Figure 2.**
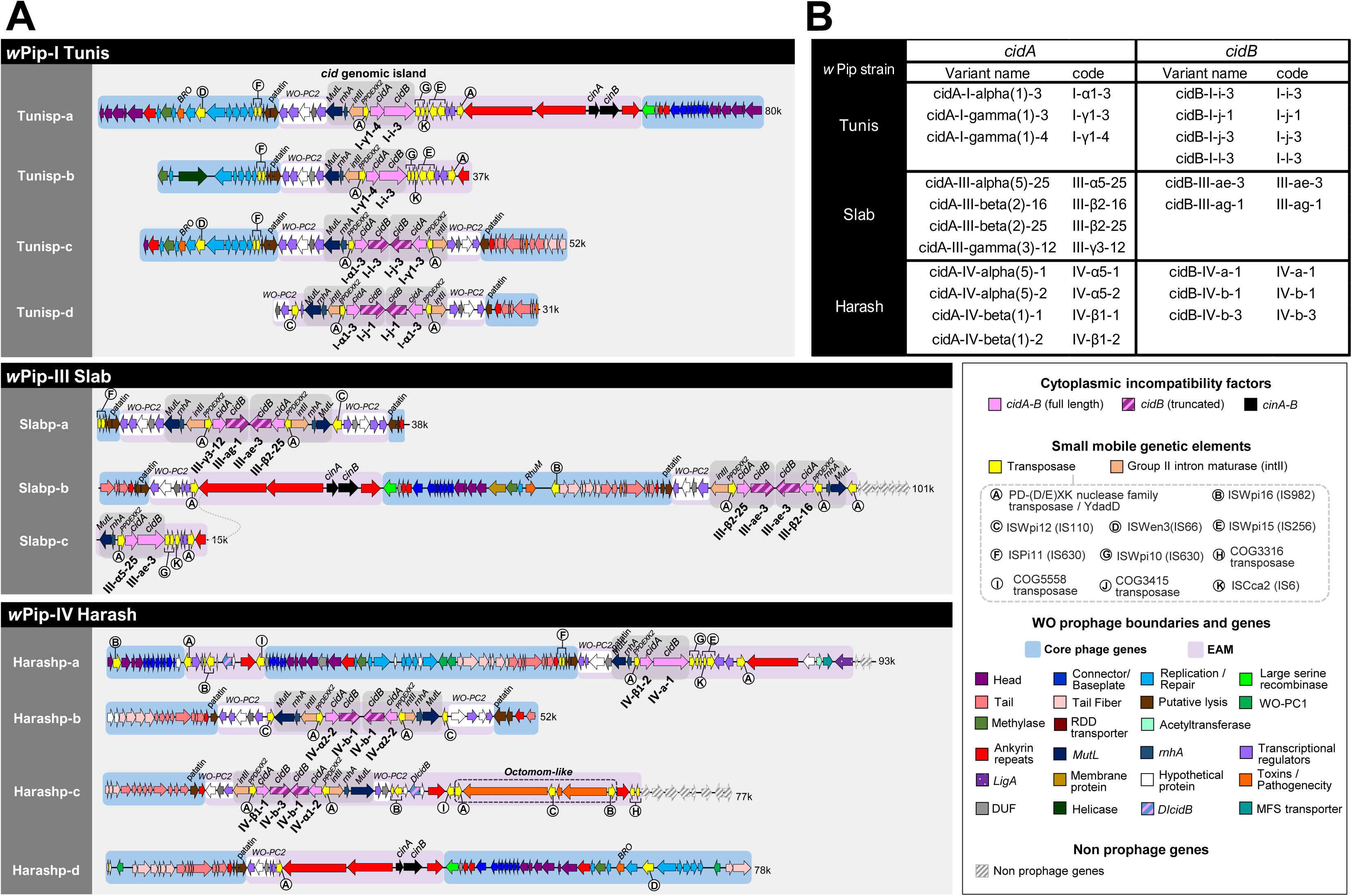
Genomic environments of the *cif* pairs in *w*Pip-I Tunis, *w*Pip-III Slab and *w*Pip-IV Harash lines. (A) Detailed gene annotations of the immediate genomic environments of the *cid* and *cin* pairs in *w*Pip-I Tunis, *w*Pip-III Slab and *w*Pip-IV Harash. The names of the different *cidA* and *cidB* variants are indicated below each corresponding gene. *DIcidB* refers to the isolated *cidB* fragments containing a deubiquitinase domain in *w*Pip-Pel (*WP1290–1291*). (B) The *cidA* and *cidB* variants previously identified in the *cid* repertoires of *w*Pip-I Tunis, *w*Pip-III Slab and *w*Pip-IV Harash lines as described by Namias et al. 2025 with help of Nanopore sequencing of PCR amplicons and their corresponding code name used in panel (A). The length of each sequence in kb is indicated at the end of each prophage region.

### All previously described *cif* repertoires were covered by our contigs

Each contig obtained from the genomes of *w*Pip-I Tunis, *w*Pip-III Slab, and *w*Pip-IV Harash includes one or two tandems of *cidA-B* variants (*cif* type I) or one tandem of monomorphic *cinA-B* (*cif* type IV). No other types of *cif* were detected. By aligning the *cif* sequences from new contigs with references from (Namias et al. 2025), the *cid* variants were identified. Such identification showed that the new contigs contained all the different *cidA* and *cidB* variants previously described in *w*Pip-I Tunis, *w*Pip-III Slab, and *w*Pip-IV Harash by Nanopore sequencing of PCR products (Namias et al. 2025) (fig 2). For example, in Tunis lineage, we successfully recovered the variants *cidA-I-alpha(1)-3, cidA-I-gamma(1)-3, cidA-I-gamma(1)-4, cidB-I-i-3, cidB-I-j-1, cidB-I-j-3 and cidB-I-l-3* (fig 2). The total number of *cid* variants found in our contigs also perfectly matched the total number of variants in these lines as investigated by qPCR (Table 1). Indeed, six *cidA-cidB* tandems were identified in the genomes of *w*Pip-I Tunis, five tandems in *w*Pip-III Slab and *w*Pip-IV Harash, while one *cinA-cinB* tandem was present in all genomes (Table 1). This suggests that our method captured all the genomic regions that contain *cif* genes in the *w*Pip genomes studied (fig 2). As previously described in Namias et al. 2025, we confirmed that only one *cidB* variant per genome was in a full length form (*i.e*. including a full DUB domain). However, the same full-length variant was found in two copies in Tunis genome, each located in distinct contigs (fig 2). All the other *cidB* variants were truncated (*i.e.* without a region including the DUB domain) and located within a palindromic organization (*cidA-cidB-cidB-cidA*) (fig 2).

### All the *cif* genes are located within WO prophage regions in *w*Pip

Annotations of the contigs demonstrated that the different *cid* variants and the monomorphic *cin* genes are all located in WO prophages (fig 2). Indeed, genomic regions flanking *cif* were always composed of (i) core phage genes (*eg.* head, tail, replication genes),(ii) WO-protein cluster 1 and 2 (WO-PC1 and WO-PC2), gene clusters with unknown function that was formalized as WO prophage markers by Bordenstein and Bordenstein 2022, and/or (iii) Eukaryotic Association Modules (EAM), that include genes with putative eukaryotic origins that are frequently described within WO prophages (Bordenstein and Bordenstein 2016; Bordenstein and Bordenstein 2022; Vancaester and Blaxter 2023; Grève et al. 2024) (fig 2). The total WO prophage regions hosting *cif* genes represent a significant part of the three *w*Pip genomes investigated here. As cumulative WO prophage regions in both *w*Pip-Pel and *w*Pip-JHB represent ∼20% of the complete genomes, and prophage regions in the new contigs containing *cif* represent 13 to 19% of the expected *w*Pip genome size (∼1.5 Mb), our data likely encompass a substantial fraction of the WO prophage regions. The phylogeny of the large serine recombinase, previously proposed as a ‘typing tool’ to discriminate clades (Sr1 to Sr4WO) within the Wovirus genus (Bordenstein and Bordenstein 2022), reveals that all WO prophages of *w*Pip strains belong to the sr3WO clade (Supplementary figures 2A and S3A). This suggests that they are phylogenetically closely related and have recently diverged. However, the analysis of other WO prophage markers reveals distinct evolutionary histories for each gene examined, with genes from *w*Pip sometimes clustering with WO prophages from other *Wolbachia* strains (Supplementary figures 2B-H and 3B-H). These analyses confirm that *w*Pip WO prophages are major recombination hotspots experiencing recurrent horizontal gene transfers.

### High diversity of WO prophages hosting *cid* genes

For each of the three phylogenetically distant *w*Pip genomes analysed here, at least three different prophage regions containing *cid* genes were recovered (fig 3). This result clearly differs from the *w*Pip-Pel reference genome, in which only the WOPip1 prophage contains a *cid* tandem (fig 1 B,C). *Wolbachia* in the Tunis isofemale line (*w*Pip-I), shows a full length *cid* version in a WOPip1-like region with well conserved gene composition (fig 3). Full length-*cid* genes in Slab and Harash (respectively in *w*Pip-III and IV groups; fig 1A) were found in mosaic prophage regions, exhibiting gene compositions which could not be assigned to any previously described WO prophage from *w*Pip-Pel (fig 3). In Tunis, Slab and Harash, the *cid* truncated palindromic variants were all identified in mosaic prophage regions (fig 3). All these prophage regions exhibit different genomic architecture (fig 3) showing a greater diversity of WO prophages in *w*Pip than previously suspected. The new annotation of transposases allowed to reveal their high diversity within WO prophages of *w*Pip (fig 1C, fig 2A). Interestingly, some *cid* variants are located in prophagic environments (Slabp-b, Harashp-a and Harashp-d) that contain the full arsenal of genes necessary to encode a complete phage particle, including head, tail, tail fiber, recombinase, putative patatin lysis, baseplate-connector, and replication-repair functions Bordenstein and Bordenstein 2022 (fig 2).

**Figure 3.**
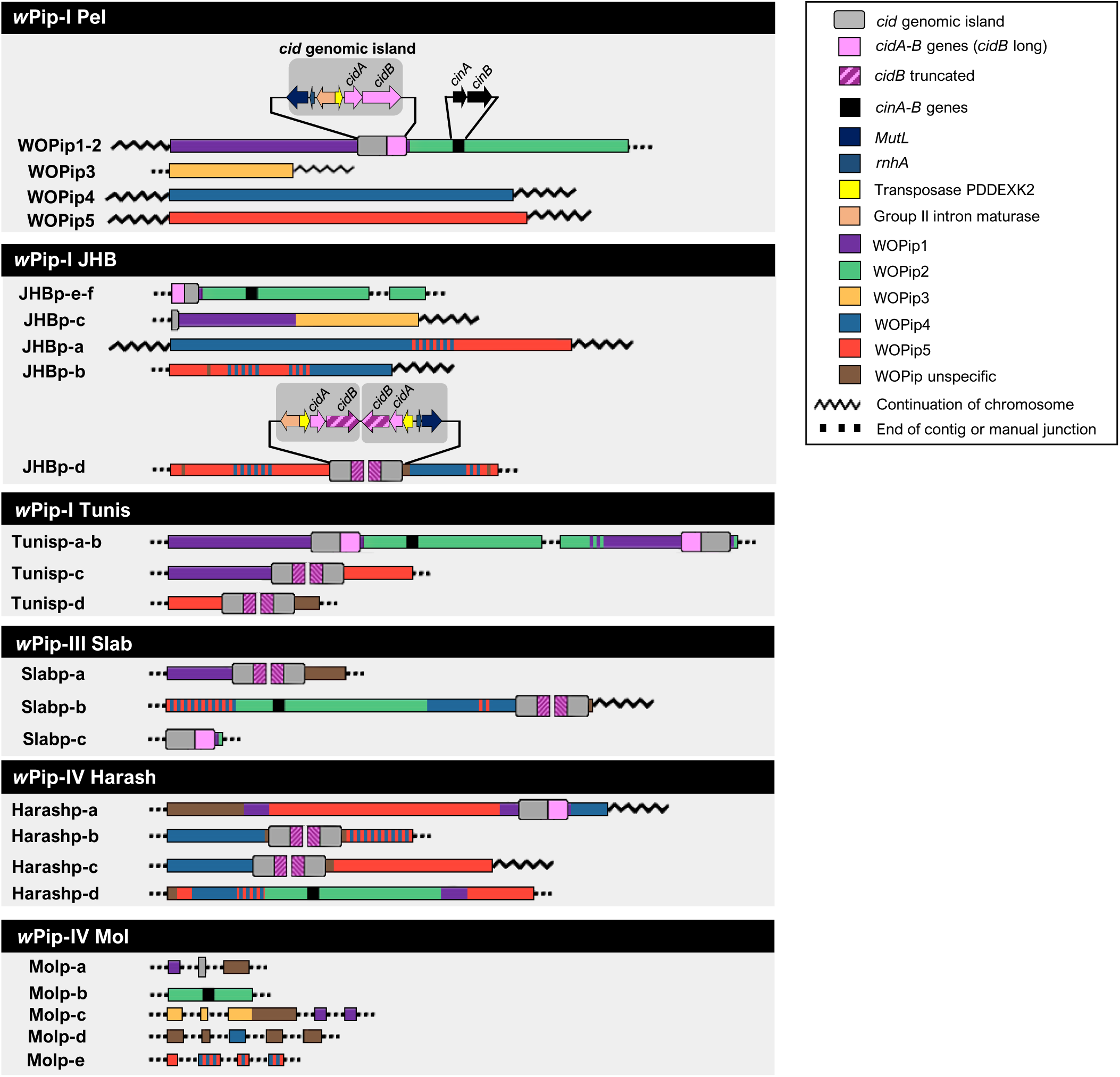
Diversity of WO prophage harboring *cif* genes among different *w*Pip genomes. This figure was built by assigning each portion of newly annotated prophage to WOPip1, WOPip2, WOPip3, WOPip4, and WOPip5 references from *w*Pip-Pel. The *cid* genomic islands and the type of *cidB* (long or truncated) are symbolized to allow their quick identification.

### Highly conserved WOPip2-like environment for *cin* genes

In all *w*Pip genomes yet studied, there is a single unique *cin* pair per genome consistently located in the same syntenic prophagic region composed of genes enriched in ankyrin repeat domains (*WP0292*, *WP0293* and *WP0296*), a large serine recombinase (*WP0297*) and a putative gene composed by domain with unknown function (DUF) (*WP0298*) (fig 1C, fig 2, fig 4A). Such prophagic environments, a part of WOPip2 in *w*Pip-Pel, are conserved both in the three genomes investigated here and previously published draft genomes (*w*Pip-Pel, *w*Pip-JHB and *w*Pip-Mol). If the *cinA* and *cinB* genes were found perfectly conserved between all the *w*Pip genomes, nucleotide polymorphism is present in some downstream *cin* neighboring genes (fig 4A).

### The “*cid* genomic islands”: hotspots of recombination for *cid* genes

#### What constitutes the “cid genomic islands”?

Four genes were found associated with *cid* among the different prophage regions annotated in this study: (i) a DNA repair gene (*MutL*, *WP0278*), (ii) a RNAse (*rnhA*, *WP0279*), (iii) a maturase required for processing of group II intron-retroelements (*intII*, *WP0280*), and (iv) a PDDEXK2 transposase composed by a PDDE(X/K) nuclease-like domain (*WP0281*) (fig. 4B, fig. 5A). These four genes and a *cid* pair form repeated genomic units that we called “*cid* genomic islands”. Indeed, these islands are present in multiple copies in distinct WO prophage regions contrary to what was previously established in the *w*Pip-Pel reference genome, where they were only present in WOPip1 (fig.1C, fig. 3).

**Figure 4.**
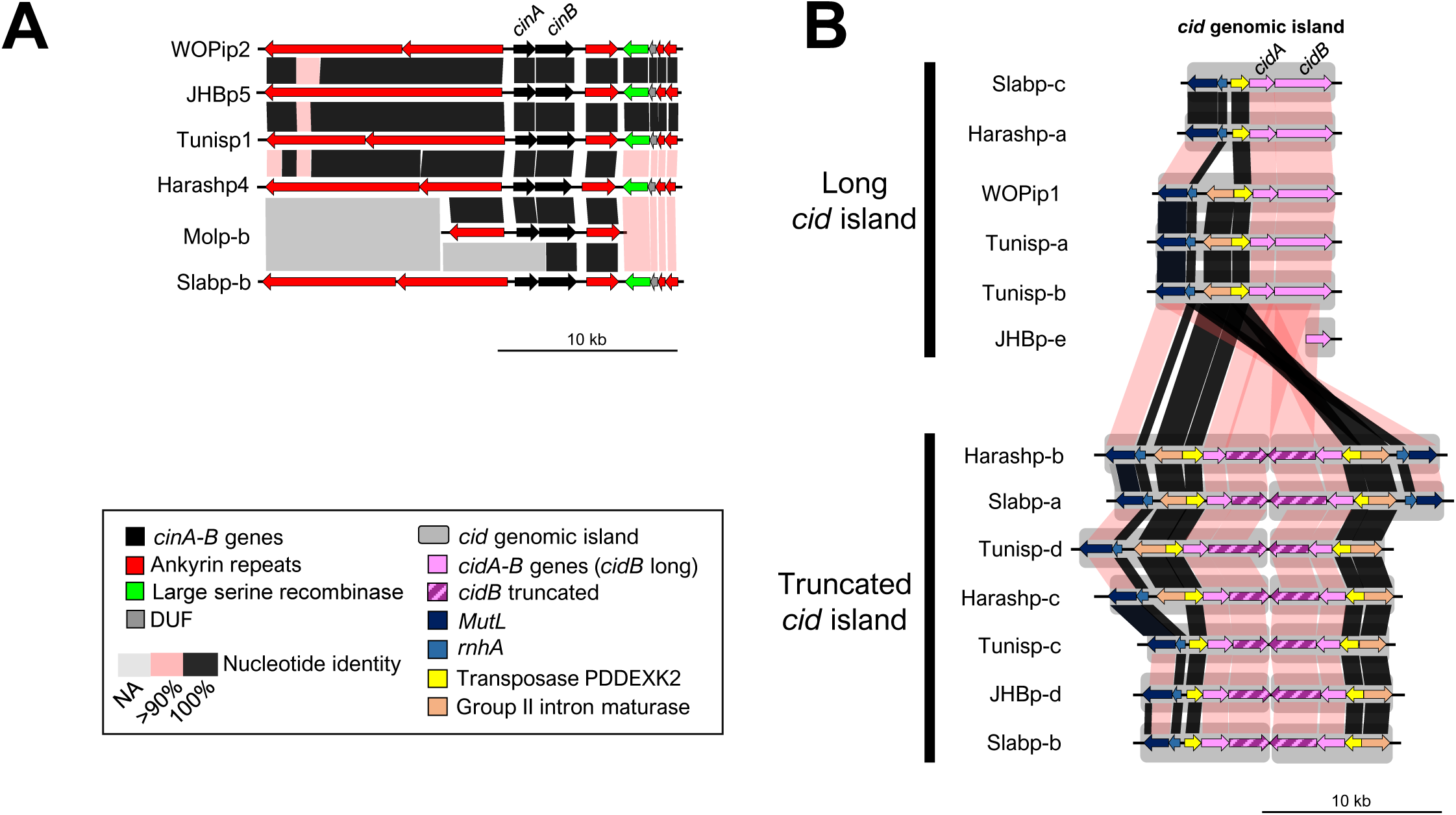
Gene nucleotide identity among genes from the (A) conserved *cin* environment and (B) among genes from the *cid* genomic islands from different *w*Pip.

**Figure 5.**
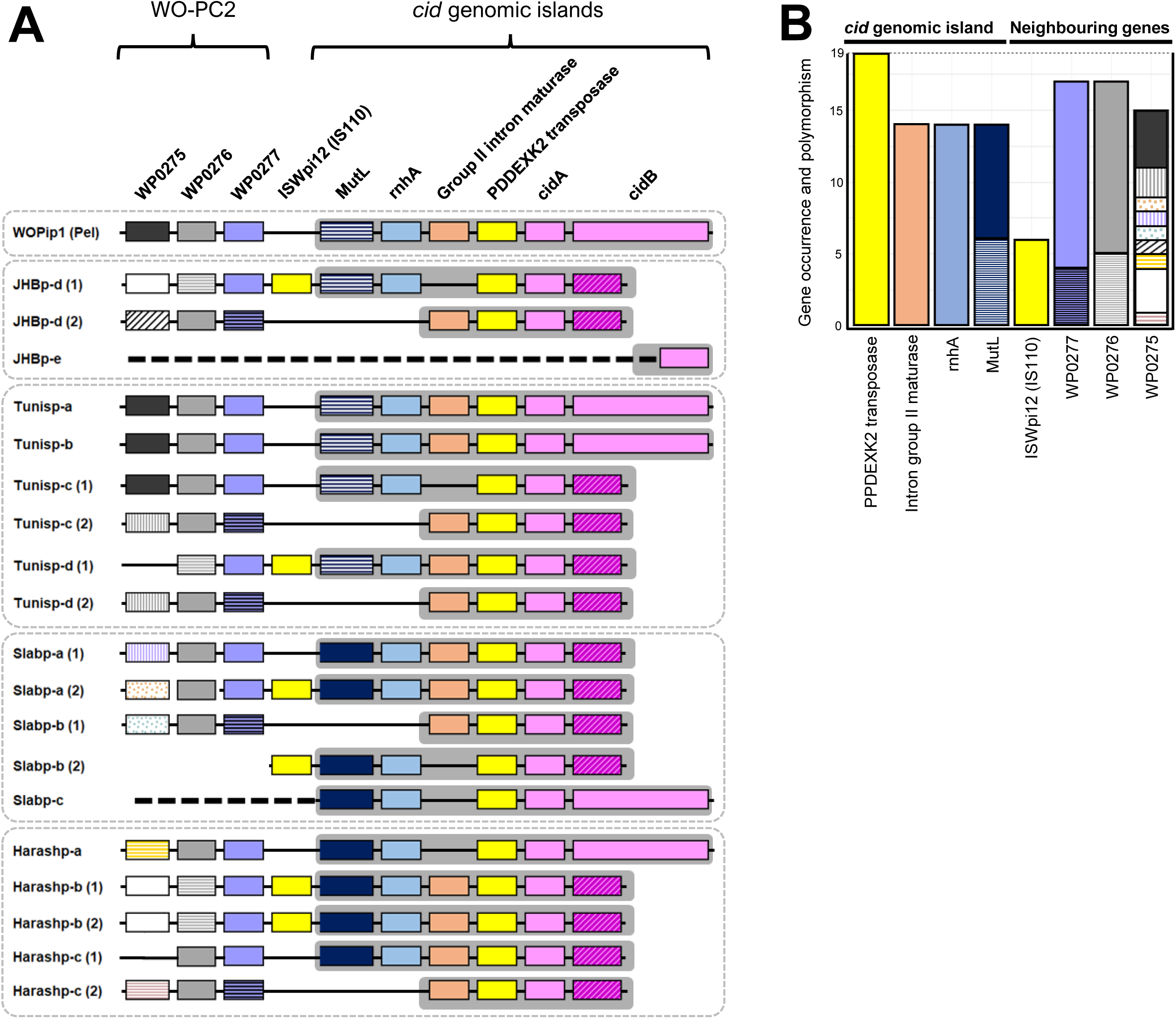
Modularity of the *w*Pip *cid* genomic islands and WO-PC2. (A) Gene compositions and variation of the 20 *w*Pip *cid* genomic islands analysed in this study. The proposed boundaries of the *w*Pip *cid* genomic islands are highlighted with a gray background. The (1) positioned after the name of some *w*Pip islands indicates that they belong to the left *w*Pip genomic island of a palindrome, while the (2) indicates the *w*Pip *cid* genomic islands that belong to the right ones. Gene codes from the *w*Pip-Pel reference genome were used to label WO-PC2 genes (*WP0275, WP0276, WP0277*). Polymorphic variants in *WP0275, WP0276, WP0277* and *MutL* are presented using different colours and motifs. Dashed black lines indicate the end of a scaffold. (B) Occurrence of different alleles in the *cid* genomic islands and WO-PC2

#### Modularity of gene composition among cid genomic islands

By comparing the gene composition of these islands, we described presence/absence patterns for certain genes, with the exception of the PDDEXK2 transposase which is the only gene always found associated with *cidA–cidB* pairs (fig. 5A-B). Otherwise, the variation in *cid* genomic islands gene composition mostly concerned the *MutL–rnhA* pair that is absent in some islands (fig 5A). The group II intron maturase is also occasionally missing (fig 5A). These recurrent patterns of gene loss and retention suggest that the *cid* genomic islands behave as modular and rearrangeable mobile elements, shaped by genomic rearrangement. At the genome scale, all *w*Pip isofemale lines (*w*Pip-I Tunis, *w*Pip-III Slab and *w*Pip-IV Harash) present different versions of *cid* genomic islands (fig 5A) but different *w*Pip lines can harbor genomic islands with the same gene composition (*e.g.* Slabp-a, Harashp-b, and Harashp-c) (fig 5A). Moreover, both full length and truncated forms of *cidB* can be found within *cid* genomic islands with identical gene composition as, for instance, in Tunis with Tunisp-a, Tunisp-b, and Tunisp-d(1) (fig 5A).

#### High nucleotide identity of genes composing cid genomic islands

Contrary to *cidA-cidB* genes that are highly polymorphic among *w*Pip lineages (Tunis, Slab and Harash) due to block recombination leading to different variants (Bonneau, Atyame, et al. 2018; Namias et al. 2024a; Namias et al. 2025) (fig 2B), the other genes of the *cid* genomic islands are highly conserved (fig 4, Supplementary Figure 4). Notably, the ubiquitous PDDEXK2 transposase (1,010 kbp), always found associated with *cid* tandems, is monomorphic among all *cid* genomic islands (fig 4, fig 5, Supplementary Figure 4A). When present, group II intron maturases (1,344 bp) and *rnhA* (558 bp) are also strictly monomorphic (fig 4, fig 5, Supplementary Figure 4B-C), as well as intergenic regions which share >99% similarity between *cid* genomic islands. For *MutL*, there is 99% identity between all islands for the first 921 bp (fig 4, fig 5, Supplementary Figure 4D). The following 1,845 bp of the gene exhibited polymorphism as previously detected and used for *w*Pip specific MLST (Atyame et al. 2011) (Supplementary Figure 4D). Depending on the versions of *cid* genomic islands, the strictly conserved sequence lengths range from approximately 2,700bp (*e.g.,* when only the group II intron maturase and PDDEXK2 transposase are present, as seen in Tunisp-c, JHBp-d, and Slabp-b) to around 5,300bp (when *MutL, rnhA*, group II intron maturase, and PDDEXK2 transposase are all present). For truncated versions of *cidB,* the most common *cidB* versions, two palindromic *cid* genomic islands face each other. Consequently, the strictly conserved sequence of the two grouped islands flanking the *cidA-cidB-cidB-cidA* genes (Namias et al. 2025), range from 3,380pb downstream (for JHBp-d, Tunisp-c and Slabp-b) to 5,306 bp flanking both sides (for Harasp-b and Slabp-a) (fig 4B). Genes from the WO-PC2 region, originally identified in prophages WOPip2, WOPip4, and WOPip5 in *w*Pip-Pel, were consistently found immediately upstream of *cid* genomic islands. Although an additional ISWpi12 element is occasionally present, WO-PC2 is conserved in gene composition across prophage regions. However, the WO-PC2 genes are highly polymorphic in nucleotide identity (fig 5B). Indeed, two variants were identified for both *WP0277* and *WP0276*, and nine variants were found for *WP0275* (fig 5B). Because of this polymorphism, we decided not to include them in the *cid* genomic islands composed of highly conserved genes.

### The flanking regions of *cid* genes in *w*Naev strengthens the hypothesis of *cid* horizontal transfer between distant *Wolbachia*

The *w*Naev genome from the lepidopteran *Rhopobota naevana* (OX366333.1) was also analyzed, as it was proven to contain multiple *cid* copies, including one *cidA-cidB* pair suspected to have laterally transferred into a *w*Pip ancestral genome (Namias et al. 2025). Five WO prophage regions were annotated containing 10 *cifA-cifB* pairs from different types including four *cid* pairs (type I) and one *cin* (type IV) (Supplementary Figure 5). Two of the annotated prophage regions, *w*Naev region 2 and region 3, harbor large serine recombinases that belong to the sr3WO clade along with *w*Pip prophages (Supplementary Figures 2A and 3A). Namias et al. 2025 hypothesized that the *cid* pair from the *w*Naev region 2, which is closely related to *cidA-cidB* pairs from *w*Pip, was the source of a horizontal transfer into a *w*Pip ancestral genome (Namias et al. 2025). We show that, in *w*Naev region 2, the *cid* pair is also flanked by two hallmark genes of *w*Pip *cid* genomic islands: the PDDEXK2 transposase and the group II intron maturase strengthening an horizontal transfer scenario of *cid* pair along with these genes (Supplementary Figure 5B). Phylogenetic analysis of these two flanking genes showed that the ones from *w*Naev region 2 perfectly cluster in the same phylogenetic clade, with respectively 99 and 100% nucleotide identity, with the ones found in *w*Pip *cid* genomic islands. This result strengthens the hypothesis of a recent horizontal transfer of a *cid* tandem along with other genes of a *cid* genomic island (Supplementary Figure 4A-B).

## Discussion

*Wolbachia w*Pip genomes contain two major types of CI Inducing Factors (*cif*): the *cid^w^*^Pip^ (also referred to as type I *cif*), whose B gene displays a DUB domain, and the *cin^w^*^Pip^ (also referred to as type IV *cif*), whose B gene encodes active nuclease sites. Transgenesis of both *cid^w^*^Pip^ and *cin^w^*^Pip^ operons in *Drosophila* can induce strong CI (Beckmann et al. 2017; Chen et al. 2019; Beckmann et al. 2019a; Sun et al. 2022). Since both *cinA* and *cinB* are completely monomorphic among *w*Pip, their involvement in the diversity of incompatibility patterns between *Culex* lineages was previously ruled out (Bonneau, Atyame, et al. 2018). On the contrary, *cid* pairs are polymorphic within and between *w*Pip genomes (Bonneau et al. 2018a; Bonneau et al. 2019; Sicard et al. 2021; Namias et al. 2024a; Namias et al. 2025). Targeted genetic approaches focusing on *cid* genes, based on Nanopore amplicon sequencing (Namias et al. 2023), have previously shown that each *w*Pip genome harbors a full length variant (with intact terminal DUB domain) and several truncated forms depleted of their terminal regions (Namias et al. 2025). The combined results from heterologous expression studies (in yeasts, drosophila cells and drosophila insects, Beckmann et al. 2017; Horard et al. 2022; Wang et al. 2022; Terretaz et al. 2023) and correlative studies in natural situation [i.e. matching *cid* diversity with *Culex* crossing experiments, Bonneau et al. 2018a; Bonneau et al. 2019; Namias et al. 2025), provide an explanation for the diversity of CI in *C. pipiens* within a toxin-antidote model (Beckmann et al. 2019b). In this model, the variability in long CidB variants (the toxins) primarily accounts for the variability in modification (*mod*) types, while variability in CidA variants (the antidotes) explains the variability in rescue (*resc*) capacities (Namias et al. 2024a; Namias et al. 2025).

While genetic studies of *cid* effectors have clarified their central role in the evolution of CI diversity in *C. pipiens*, the genomic context underlying their amplification and diversification had yet to be explored. In this study, we focused on three *w*Pip lineages that induce distinct crossing compatibility patterns and whose *cid* repertoires have already been fully characterized (Supplementary Table 1, Atyame et al. 2014; Sicard et al. 2021; Namias et al. 2025). Using long-read sequencing, we reconstructed the genomic regions flanking each previously characterized *cid* variant, resulting in robustly assembled contigs. Annotation of these contigs enabled us to investigate the genomic drivers underlying *cid* amplification and diversification. Moreover, such data allowed us to question the difference in polymorphism between the two types of *cif* effectors in *w*Pip, *cid^w^*^Pip^ and *cin^w^*^Pip^, which are present in all genomes but exhibit distinct molecular evolution patterns: *cinA-B^w^*^Pip^ is found in a single monomorphic copy versus *cidA-B^w^*^Pip^ in multiple (up to 6 different) recombinant copies per genome.

### All *cif* genes are located in WO prophages challenging the assembly of *w*Pip

The contigs obtained allowed us to cover all variants of *cid* genes (i.e. the full *cid* repertoire) as well as the monomorphic pair of *cin* genes for each different genome (fig 2). All these *cif* genes were found in proximity to core phage genes and/or genes of the EAM, both defined as hallmarks of WO prophages (Bordenstein and Bordenstein 2022). All the prophage regions containing *cif* genes in *w*Pip belong to Wovirus genus, more specifically to the sr3WO clade, based on the phylogeny of their large serine recombinases (Bordenstein and Bordenstein 2022) (Supplementary Figures 2A and 3A). Monomorphic *cin* genes are embedded in highly syntenic prophage regions in all analysed *w*Pip genomes, covering over nearly 36,000 bp involving almost all genes constituting WOPip2 described in the reference *w*Pip-Pel genome (Klasson et al. 2008; Bordenstein and Bordenstein 2022). In contrast, most of the *cid* variants are located in variable WO prophages that differ in gene compositions within and between *w*Pip lines which are mosaics of the reference WO prophages described in the *w*Pip-Pel reference genome (Klasson et al. 2008). Altogether, these results indicate that WO prophage are highly dynamic across *w*Pip lineages, as previously suggested by Reveillaud et al. 2019. Their complex repetitive architecture not only poses a major challenge for genome assembly but also makes it impossible that a single reference genome fully represents their diversity. Our study provides new insights into this difficulty by revealing that WO prophages harbouring *cid* genes exhibit highly variable gene compositions, yet consistently containing multiple repeated core genes (*e.g*. baseplate, head, tail) and many copies of the WO-PC2 elements, which likely hinder genome circularization during assembly. The direct examination of Nanopore long reads polished with Illumina data, combined with a prior, precise identification of the expected *cid* gene repertoires for each sequenced *w*Pip lineage (Namias et al. 2025), proves to be an effective strategy for circumventing the issue of reconstructing WO prophages. The cumulative size of WO prophage regions has been identified as a major driver of genome size variation in *Wolbachia*, as shown by the strong correlation observed across 78 supergroup B genomes (Vancaester and Blaxter 2023). Using this correlation, with a predicted full genome size of ∼1.5 Mb (Klasson et al. 2008), approximately 20% of the *w*Pip genomes are expected to be occupied by WO prophages (Vancaester and Blaxter 2023). In *w*Pip-I Tunis, *w*Pip-III Slab, and *w*Pip-IV Harash, we found that WO prophage regions harboring *cif*, represent up to 19% of the complete genome. These results suggest that most WO prophages in *w*Pip might harbor *cif* genes.

### *cid* genes are part of a modular and mobile genomic island

The presence of WO core genes along with WO-PC2 in the close genomic environments of all *w*Pip *cid* pairs proved that they belong to prophagic regions. However, the gene compositions of these WO prophages are highly variable suggesting important structural genomic rearrangement in these regions (fig 3). The phylogenetic analysis of the *w*Pip WO prophage core genes showed that each of them has its own evolutionary history suggesting recurring horizontal transfer and recombination events (Supplementary figures 2 and 3). Unexpectedly, in these highly dynamic prophagic environments, we show that *cid* genes are tightly associated with four modular genes that exhibit high sequence conservation. We propose that these genes form, along with *cid* genes, genomic islands that move between different WO prophage regions. Among the four non-*cid* genes forming the “*cid* genomic islands”, two are mobile elements, and the other two appear to be implicated in recombination and DNA repair. The first mobile element, a PDDEXK2 transposase (*WP0281*) consistently found just upstream of *cidA* in *w*Pip genomes, was already described as broadly associated with *cif* genes in *Wolbachia* (Tan et al. 2024; Amoros et al. 2025). The PD-(D/E)XK domain from the PDDEXK2 transposase shares the same name as nucleases found in all the *cifB* types (LePage et al. 2017; Beckmann et al. 2017; Martinez et al. 2021), although showing no alignable sequence similarity. Transposases such as Insertion Sequences (IS) had already been proposed as potential drivers of horizontal transfers of *cif* genes in other *Wolbachia* and bacterial endosymbionts (Cooper et al. 2019; Madhav et al. 2020; Oswalt et al. 2025; Amoros et al. 2025). These transfers, involving *cif* genes flanked by IS elements, can occur within distinct WO prophage regions, as observed in the *Wolbachia* endosymbiont of the horn fly *Haematobia irritans irritans*, *w*Irr (Madhav et al. 2020). The second mobile element found associated with *cid* genes in *w*Pip is the Group II Intron maturase (intII), a type of transposable element with a reverse transcriptase activity previously reported in several *Wolbachia* strains (Leclercq et al. 2011), and sometimes also found in association with *cif* genes in *Wolbachia* and other bacterial endosymbionts (Bordenstein and Bordenstein 2022; Tan et al. 2024; Amoros et al. 2025). Apart from mobile elements, two other genes are strongly associated with *cid* genes in their genomic islands: (i) the *rnha* gene, which encodes an RNase H enzyme that may play a role in RNA regulation, and (ii) *MutL*, a gene involved in the DNA mismatch repair system. *MutL* was punctually found near *cif* genes in other genomes such as *w*AlbB (inside the WO-like island WOAlbB3) or in the *Candidatus* Mesenet longicola, a bacterium phylogenetically close to *Wolbachia* (Takano et al. 2021; Bordenstein and Bordenstein 2022; Amoros et al. 2025). While primarily known for its role in DNA mismatch repair, the recurrent presence of *MutL* in *w*Pip *cid* genomic islands suggests a role in the maintenance, regulation, or even remodelling of these islands. Its activity could contribute to preserving the integrity of horizontally transferred genes, modulating local mutation rates, or facilitating recombination events, thereby supporting both the stability and adaptive diversification of *cid* loci within and across *Wolbachia* genomes. The *MutL* found in the *cid* genomic islands has been proposed to have a phage-related origin, as it is divergent from chromosomally encoded *MutL* orthologs (Fallon 2021). The *rnhA* gene remains also functionally uncharacterized in *Wobachia*. However, its known role in resolving RNA–DNA hybrids (Schroeder et al. 2023) suggests a potential involvement in transcriptional regulation, maintenance of genome stability, or modulation of recombination within the dynamic genomic environment of *cid* genomic islands. Although the PDDXK2 transposase, intII, and *MutL* have been previously identified separately as frequently associated with *cif* genes, they have never been reported as being jointly associated within a single conserved unit alongside *cif* genes. Altogether, this configuration forms a mobile, identifiable unit that is both stable and recombinogenic, making *w*Pip *cid* genomic island a unique case for *cif* genes. Interestingly, the *w*Pip *cid* genomic islands share architectural similarities with another genomic island, the Octomom module known for its strong instability in *w*Melpop (Chrostek et al. 2013; Chrostek and Teixeira 2015; Chrostek and Teixeira 2018; Duarte et al. 2021; Monnin et al. 2021; Bénard et al. 2021). The Octomom module is a genetic element whose copy number can evolve rapidly (increase or decrease over a few generations), and which is directly correlated with *Wolbachia* density within the host (Chrostek et al. 2013; Chrostek and Teixeira 2015; Chrostek and Teixeira 2018; Duarte et al. 2021; Monnin et al. 2021; Bénard et al. 2021). Similar to the *cid* genomic islands in *w*Pip, the Octomom module in *w*MelCS and *w*Melpop is flanked by a PDDEXK2 transposase, two reverse transcriptases (showing homology with intII), and a *MutL* gene (Bordenstein and Bordenstein 2022). This shared genetic architecture between *cid* genomic islands in *w*Pip and Octomom modules in *w*Mel suggests that both may rely on a common set of mobility and regulatory genes to facilitate their amplification, structural rearrangement, and/or stability.

### High nucleotide identity of the genomic islands increases recombination probabilities during DNA replication

Although the gene composition of the *cif* genomic islands is modulable (fig 4 and 5), the four genes show high sequence similarities (>99%) between islands. Thus, upstream of each *cidA-cidB* pair, there are between 2,700 bp and 5,300 bp that are strictly identical between two islands within the same genome or between genomes of different *w*Pip lineages. Some palindromic structures harboring truncated *cid* pairs are flanked by identical sequences totalling up to 10,600 bp. The presence of highly conserved regions near *cid* pairs (especially in a palindromic manner) increases the probability of recombination (homologous or gene conversion) between genomic regions flanked by these repetitions (Rocha 2003; Vos 2009; Paulsson et al. 2017). High recombination frequency may explain: (i) why the number of *cid* pairs increases or decreases between *w*Pip strains and over the evolution of a single isofemale lineage (Namias et al. 2024a); and (ii) why there is a high probability of recombination between *cid* genes, which has been shown to be a major driver of their diversification (Namias et al. 2025). Among the available genomes, another *Wolbachia* strain, infecting the lepidopteran *Rhopobota naevana*, so-called *w*Naev, also exhibits the pattern of *cid* amplification with all the five prophagic regions harboring one *cif* tandem (Supplementary Figure 5). It has been proposed that the formation of palindromic structures involving truncated *cidB* genes in *w*Pip may have resulted from a horizontal transfer event between a *w*Naev-like strain and an ancestral *w*Pip strain (Namias et al. 2025). Our genomic data shows that even if prophage regions from *w*Naev and *w*Pip both belong to srWO3 clade based on large serine recombinase, other WO prophage markers are phylogenetically discordant ruling out the hypothesis of a full WO phage exchange between the ancestral genomes of these two strains (Supplementary Figure 2). However, in the prophagic regions in *w*Naev, the *cid* tandems are also associated with genes found in *w*Pip *cid* genomic islands including the PDDEXK2 transposase and the intron group II maturase which exhibit 99% similarities with those of *w*Pip (Supplementary Figure 4). These results suggest that *cid* genomic islands could have been a main driver of horizontal transfer between these two ancestral strains.

### Genomic islands as drivers of *cid* amplification and diversification in *w*Pip

*Wolbachia* represent a distinctive group of endosymbiotic bacteria in that they are heavily infected by bacteriophages, unlike their closest relatives among the Rickettsiales (Darby et al. 2007). The high prevalence of *cif* genes in *Wolbachia* compared to other Rickettsiales may be linked to WO prophages, which constitute the predominant genomic environment of *cif* and are thought to facilitate horizontal transfer (Martinez et al. 2021; Vancaester and Blaxter 2023; Amoros et al. 2025). Previous studies have proposed that these bacteriophages, by alternating between prophagic and particle forms, could mediate the horizontal transmission of key genes between distantly related *Wolbachia* strains co-infecting the same arthropod cells (Bordenstein and Wernegreen 2004; Bordenstein and Reznikoff 2005; Bordenstein and Bordenstein 2016; Perlmutter et al. 2019; Bordenstein and Bordenstein 2022). However, sequencing of WO phage particles remains limited, and none have been shown to encapsidate *cif* genes to date (Bordenstein and Bordenstein 2016; Kupritz et al. 2021). In *w*Pip, particle-like structures resembling WO phages have been observed (Sanogo and Dobson 2006), yet no genomic evidence has confirmed that these particles are indeed WO phages that might carry *cid* genes. This absence of direct evidence suggests that transduction may not be the principal mechanism underlying the amplification and diversification of *cid* genes, and that alternative processes are likely involved. In this sense, the modularity and rearrangeability natures of the *cid* genomic islands could promote recombination during DNA replication, providing a robust mechanism for their rapid diversification. Outside *Wolbachia*, genomic islands play a crucial role in the evolution of diverse bacterial species by spreading accessory genes, including those related to antibiotic resistance, defense systems, virulence, and catabolic functions, ultimately enabling the emergence of new metabolic pathways (Juhas et al. 2009; Li and Wang 2021; Wang and Dagan 2024; Krishnakant Kushwaha et al. 2024). By rapidly supplying new adaptive traits, they also favor the local and fast adaptation of their bacterial hosts to changing environments (Juhas et al. 2009; Bardaji et al. 2017; Li and Wang 2021; Krishnakant Kushwaha et al. 2024). Here, we provide evidence that such genomic islands may also promote the diversification of CI genes in *Wolbachia*.

## Conclusion

By exploring the genomic environments of *cif^w^*^Pip^ genes, our study unveils new insights on the microevolution of highly related *Wolbachia* genomes. Our investigation of the genomic environments of *cif* types in *w*Pip reveals a strong link between their genetic diversity and the genomic architecture of the prophage regions in which they are embedded. The *cin^w^*^Pip^ genes, which are monomorphic and not involved in the diversity of CI patterns in *Culex*, are located within a conserved prophage region across all available genomic data. In contrast, *cid^w^*^Pip^ genes are embedded within conserved genomic islands but in highly variable WO prophages both within and between *w*Pip strains. These genomic islands appear to promote the dispersion and diversification of *cid* genes across genomes. This genomic dynamic likely underlies the extensive diversity of CI patterns observed in *C. pipiens*, which can evolve rapidly over just a few generations, as it alters the repertoires of toxins (CidB) and antidotes (CidA) (Namias et al. 2024a). Therefore, the evolution of CI patterns in *C. pipiens* would be mainly driven by mobile elements within WO prophage regions. Identification of the drivers of *Wolbachia* genomic evolution is of primordial importance to develop safe and sustainable *Wolbachia*-based antivectorial methods. The ability of WO prophage regions to rapidly diversify CI genes suggests that *Wolbachia* strains introduced into mosquito populations might change over evolutionary timescales, potentially affecting the stability and efficacy of vector control.

## Data availability

The WO prophage contigs reconstructed in this study for *w*Pip-I Tunis, *w*Pip-III Slab and wPip-IV Harash were deposited in National Center for Biotechnology Information (NCBI) under the accession number XXX-XXX. The Illumina polished Nanopore-sequences selected for this study are available on https://doi.org/10.5281/zenodo.17245713.

## Acknowledgments

The authors are grateful to Sandra Unal and Patrick Makoundou for technical support and maintenance of the mosquito colonies. We also thank Brandon S. Cooper and William Conner for the Nanopore sequencing of the Tunis and Harash lines, as well as Richard Cordaux for the discussion and guidance. The authors also acknowledge the ISO 9001 certified IRD i-Trop HPC (South Green Platform, www.bioinfo.ird.fr; www.southgreen.fr) at IRD Montpellier for providing HPC resources that have contributed to the research results reported within this paper.

## Author contributions

Conceptualization: J.A, A.N., M.W., M.S.; Funding acquisition: M.W., M.S, H.A.; Investigation: J.A, A.N, T.D, H.A., J.M., M.S.; Data curation: J.A, A.N, T.D, H.A., J.M., M.S.; Formal analysis: J.A.; Methodology: J.A., M.S; Supervision: M.S.; Visualization: J.A., Writing – original draft: J.A., M.S.; Writing – review & editing: All authors; Validation: M.S.

---

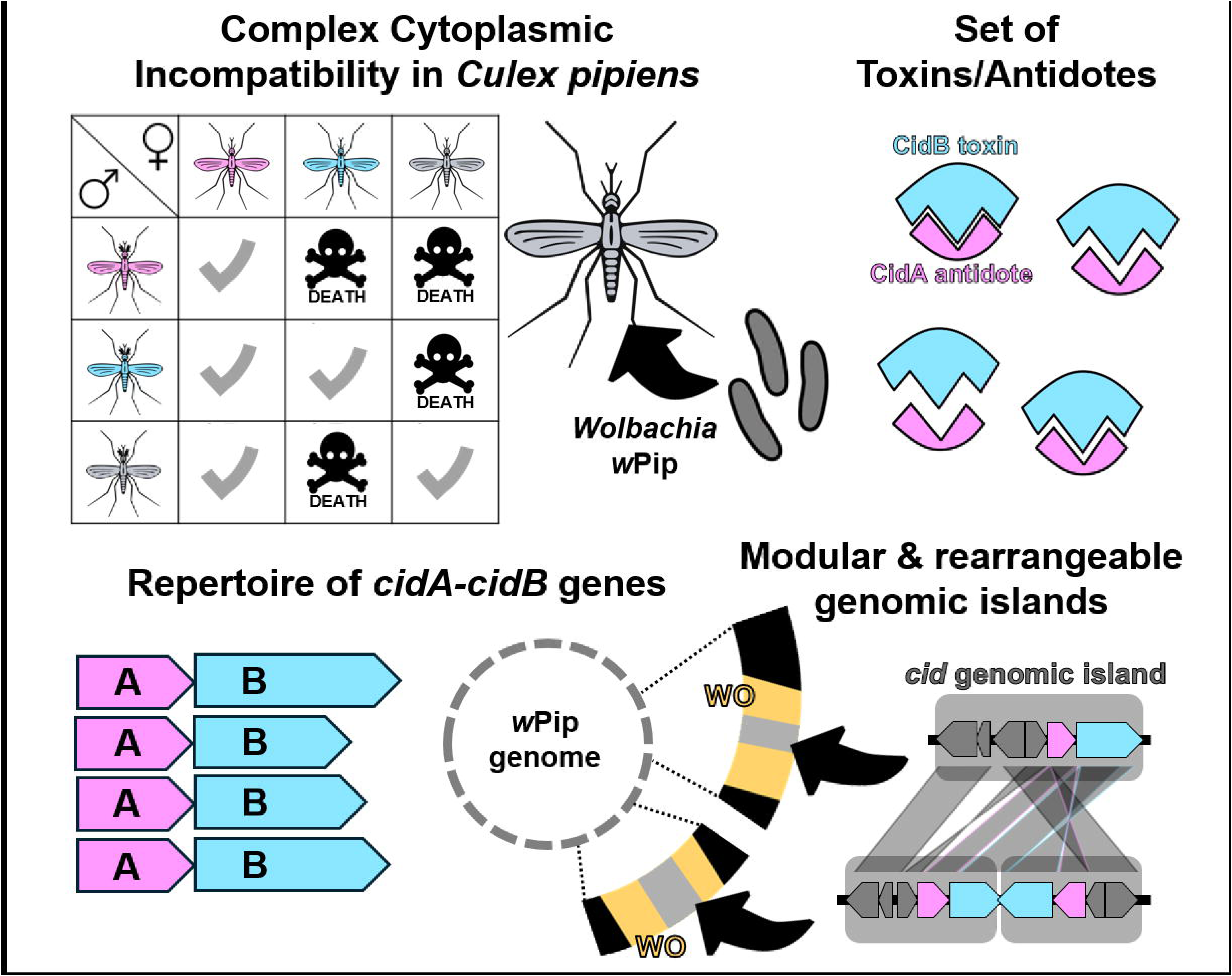

